# From Single Molecular Detail to Subcellular Dynamics: Real-Time Kinetics Study of dPA Uptake with Raman-based Spectroscopy

**DOI:** 10.1101/2025.11.21.689687

**Authors:** Aleksandra Borek-Dorosz, Patrycja Dawiec, Kenta Temma, Heqi Xi, Itsuki Yamamoto, Krzysztof Brzozowski, Joanna Profic-Paczkowska, Menglu Li, Magdalena Cudak, Kacper Siąkała, Jing Qiao, Katarzyna Majzner, Katsumasa Fujita, Malgorzata Baranska, Anna Maria Nowakowska

**Affiliations:** Jagiellonian University in Kraków, Faculty of Chemistry, Department of Chemical Physics, 2, Gronostajowa St., Krakow, Poland; Jagiellonian University in Kraków, Doctoral School of Exact and Natural Sciences, 11 Lojasiewicza St., Krakow, Poland; The University of Osaka, Department of Applied Physics, 2-1 Yamadaoka, Suita, Osaka, Japan

**Keywords:** deuterated palmitic acid, Raman spectroscopy, real-time tracking of metabolism, kinetic analysis of uptake, 3D spectroscopic imaging

## Abstract

Real-time observation of cellular transformations at the subcellular level provides insight into the dynamic biochemical processes underlying cell function and disease. However, many challenges remain in developing methods to analyze metabolic transformations in cells quantitatively. Our study applies Raman spectroscopy (RS) and stimulated Raman scattering (SRS) to link single-molecule resolution with cellular-scale lipid turnover tracking. Using HL-60 cells as a model, we monitor the uptake kinetics of deuterated palmitic acid (dPA), a marker of *de novo* lipogenesis, from the first minutes to hours.

SRS enables microscopic, high-temporal-resolution visualization of dPA incorporation across entire cell volumes, capturing both its distribution and uptake rate. Complementary RS measurements identify distinct metabolic phases: rapid incorporation into pre-existing lipid droplets (LDs) followed by the emergence of new, partially unsaturated LDs associated with cytochrome-linked metabolism. Statistical and spectral analyses reveal cell-to-cell heterogeneity, emphasizing the complexity of lipid metabolism at both the cellular and subcellular levels.

By integrating SRS and RS data into time-dependent kinetic profiles, we establish a unified submicron-to-cellular model of fatty acid metabolism. This multi-resolution spectroscopic approach demonstrates how real-time, label-based tracking can uncover dynamic metabolic heterogeneity, advancing understanding of the fatty acid uptake and metabolism in the microscale.

## 1. Introduction

Lipids comprise a diverse group of active biomolecules serving as membrane components ^[1]^, energy stores in lipid droplets (LDs) ^[2]^, and cellular mediators *via* receptor-mediated pathways ^[3]^. Their highly dynamic metabolism involves lipid uptake, *de novo* synthesis, transport, and degradation ^[4]^. In the body, these processes fluctuate throughout the day and can be influenced by a wide range of factors, including circadian rhythm, physical activity, and food intake ^[5]^. Even at the level of individual cells, LDs are not static organelles but highly dynamic structures that act as vigorous hubs of cellular metabolism, whose spatiotemporal heterogeneity is essential for balancing the deposition and release of distinct lipid species^[6]^.

Alterations in lipid metabolism, especially the accumulation of LDs, are recognized as a hallmark of inflammation ^[7,8]^, malignant progression ^[9]^, atherosclerosis ^[10]^, diabetes ^[11,12]^, or Alzheimer’s disease ^[13,14]^. Numerous studies demonstrate that fatty acid (FA) biosynthesis and oxidation are altered during tumorigenesis ^[15,16]^. Compared with normal cells, FA synthase activity increases significantly in cancer cells ^[9,15,17,18]^, including leukemias [20], and is closely associated with tumor aggressiveness and resistance to treatment ^[9,19]^.

Fluorescence techniques have undergone significant evolution, transitioning from simple lipid stains to sophisticated live-cell and flow cytometric assays capable of quantifying fatty acid transport dynamics ^[20–22]^. Nevertheless, accurately assessing fatty acid uptake remains challenging, as fluorescence- and cytometry-based assays depend on external labels that can alter cellular physiology, are prone to photobleaching, which affects signal intensity, and they also lack molecular resolution ^[22]^. On the contrary, Raman-based techniques ^[23]^ provide molecular-level information on lipid composition, localization, and dynamics ^[8,24–29]^. Recent studies using CARS and SRS microscopy have tracked FAs uptake and turnover across various cell types, revealing lipid-metabolism heterogeneity. For example, Mitra *et al*. analyzed lipid influx and turnover kinetics in human metastatic prostate cancer cells using CARS ^[30]^. Integration of Raman imaging with labeled counterparts of FAs was also used by Stiebing *et al.* to study how human macrophages store exogenous FAs over extended incubation periods ^[31]^ or by Ranneva *et al.* to analyze concentration-dependent deuterated FA (dFA) uptake quantitatively ^[32]^. Further, the combination of nonlinear optical microscopy and Michaelis–Menten kinetic modeling enabled quantitative assessment of FAs turnover, uncovering metabolic adaptations characteristic of T47D breast cancer cells ^[33]^. On the other hand, Zhang et al. proposed a method focused on the analysis of LDs displacement and velocity to distinguish metabolic states ^[34]^. The presented examples demonstrate that spectroscopy offers significant potential for quantitative kinetic analysis of FAs dynamics. However, much remains to be done, including systematic standardized kinetic analyses across different temporal regimes, particularly during the initial stages of uptake, as well as investigations at both the subcellular and whole-cell levels. Equally important is statistical analysis to capture the heterogeneity of these processes. In the future, such studies may enable the use of kinetic profiles as an additional diagnostic and prognostic tool for patients.

Given the pressing need for new methods to track the dynamics of lipid transformations in cells over time, we propose a spectroscopy-based approach to study the spatiotemporal kinetics of dPA uptake at the subcellular level and to track intracellular heterogeneity at the single-LD level. The proposed method enables the characterization of metabolism kinetics across different time regimes: short with 45-second intervals and long at ~15-minute or ~25-minute intervals. We also aimed to analyze cell-to-cell variability in dPA uptake across a large cell population and to relate the observed diversity to the molecular heterogeneity by analyzing the spectral fingerprint of cells incubated with dPA.

## 2. Results

### 2.1. DPA is a broadly applicable molecular isotopic tracer that enables monitoring the uptake of exogenous lipids into individual LDs

To verify the effectiveness of dPA as a molecular tracer in studies utilizing Raman-based spectroscopies, we compared the morphology (**Figure 1A**) and lipid composition of cells incubated with native PA and dPA with control fixed cells (3D: **Figure 1A-C**, 2D: **Figure 1D**). The use of dPA allowed us to selectively map the incorporation of exogenous FAs into single LDs using RS (**Figure 1C&D**). This is primarily possible due to the presence of distinct C–D vibrational bands in the silent region of the Raman spectrum, which are absent in endogenous molecules and uniquely associated with dPA (**Figure 1F**). Notably, high-resolution RS imaging revealed chemical heterogeneity within individual LDs in cells exposed to dPA, distinguishing domains enriched in exogenous versus endogenous lipids (**Figure 1D**). Such separation was absent in PA-treated cells, where all lipids appeared as a single undifferentiated pool (**Figure 1D**). These findings highlight dPA as a selective tracer for tracking exogenous lipid uptake and partitioning within cellular lipid stores.

**Figure 1.**
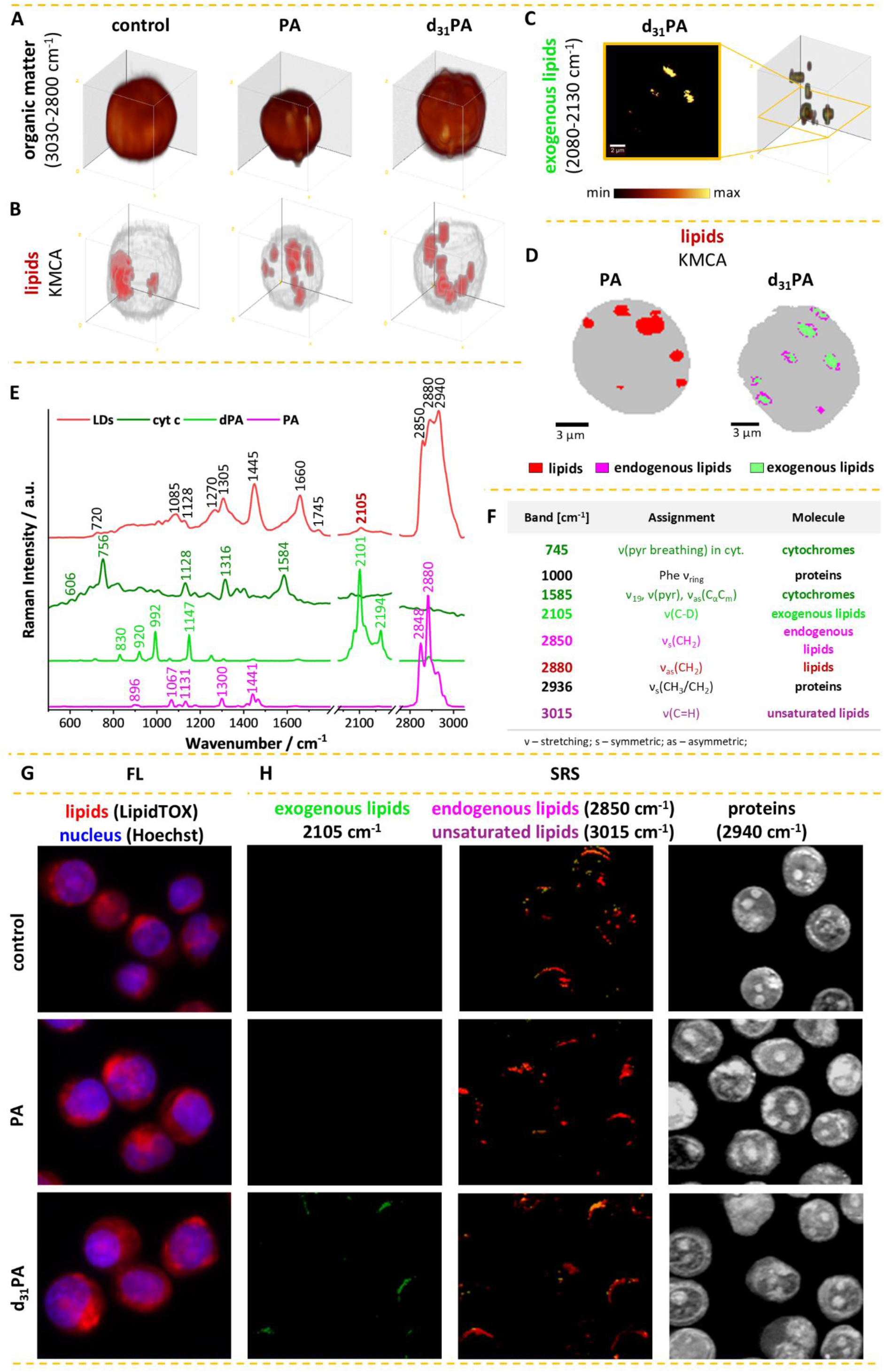
DPA as an isotopic tracer that enables the assessment of exogenous lipid uptake and incorporation within single LDs. 3D visualization of cellular morphology (integration in 2800-3030 cm^−1^ range) (A) and all lipids (red - lipids, grey – cytoplasm and nucleus) (B) in control cells and cells incubated with PA and dPA compared with 3D visualization of exogenous lipids in dPA-treated cells (integration in 2080-2130 cm^−1^ range) (C). 2D HR images showing the separation of exo- and endogenous lipids in cells incubated with dPA in contrast to cells treated with native PA, showing a single lipid pool (D). Exemplary mean spectrum of LDs compared with the spectra of reference compounds, including cyt. c, PA, and dPA (E) with marker bands characteristic of these molecules (F). Fluorescence images (Hoechst 3342 nuclei, blue and LipidTOX lipids, red) of examined cells (magnification 40x, NA=0.55) (G) compared with SRS images collected for bands specific to exogenous lipids (2105 cm^−1^, green), endogenous lipids (2850 cm^−1^, magenta), and proteins (2940 cm^−1^, grey).

Reference spectra of selected compounds, including cytochrome c (cyt. c), dPA, and native PA, were recorded to verify spectral marker positions compared to the mean spectrum of LDs of cells treated with dPA (**Figure 1E**). This approach allowed us to identify a set of characteristic marker bands, presented in a table shown in **Figure 1F**. The use of proposed marker bands facilitated selective tracking of exogenous fatty acid incorporation at the level of individual LDs (2105 cm^−1^), as well as enabled monitoring of additional molecular transformations, including cytochrome-related protein changes (745, 1585 cm^−1^), the presence of unsaturated lipids (3015 cm^−1^), and the distinction between exogenous and endogenous (2850 cm^−1^) lipid species in a lipid poll (2880 cm^−1^) which will be discussed in detail in the following chapters of the article.

Results obtained with RS were compared with those from SRS and FL microscopy (**Figure 1H&G**). Using FL microscopy with LipidTOX staining, we monitored neutral lipid accumulation in cells. Complementarily, using SRS, we visualized LDs formation and their molecular composition in response to incubation with either native PA or dPA. By targeting four characteristic Raman bands, we selectively tracked exogenous lipids (2105 cm^−1^), endogenous lipids (2850 cm^−1^), unsaturated lipids (3015 cm^−1^), and proteins (2940 cm^−1^), providing multiplexed chemical imaging at the subcellular level. Our results show that FL, as well as non-labeled RS and SRS, did not allow for the differentiation of pools of exogenous and endogenous lipids in LDs. These results demonstrate that combining coherent and non-coherent Raman spectroscopy with isotopic labeling enables selective tracking of exogenous FA uptake and incorporation into individual LDs at the submicron scale.

### 2.2. Spatiotemporal monitoring of cellular metabolism inside single cells

In our study, we employed RS to monitor the spatiotemporal dynamics of selected biomolecules, using previously defined characteristic spectral bands (**Figure 1F**). These included proteins (1000, 2940 cm^−1^), cytochromes (745 and 1585 cm^−1^), lipids (2880 cm^−1^), both endogenous (2850 cm^−1^) and exogenous (2105 cm^−1^), and unsaturated lipids (3015 cm^−1^) (**Figure 2B**). **Figure 2A** shows a schematic representation of the experimental workflow, including dPA labeling and Raman-based imaging. The experiments involved adding dPA and monitoring biochemical changes in live cells at successive time points. Results are presented for a representative live HL-60 cell, demonstrating that the time-dependent dynamics of six distinct biomolecular species can be simultaneously tracked at the subcellular level in real time (**Figure 2B**). In **Figure 2B**, these relations are represented using band-specific integration maps. At the same time, a complementary MCR analysis was performed, with the results provided in the SI, confirming the results presented in **Figure 2A**.

**Figure 2.**
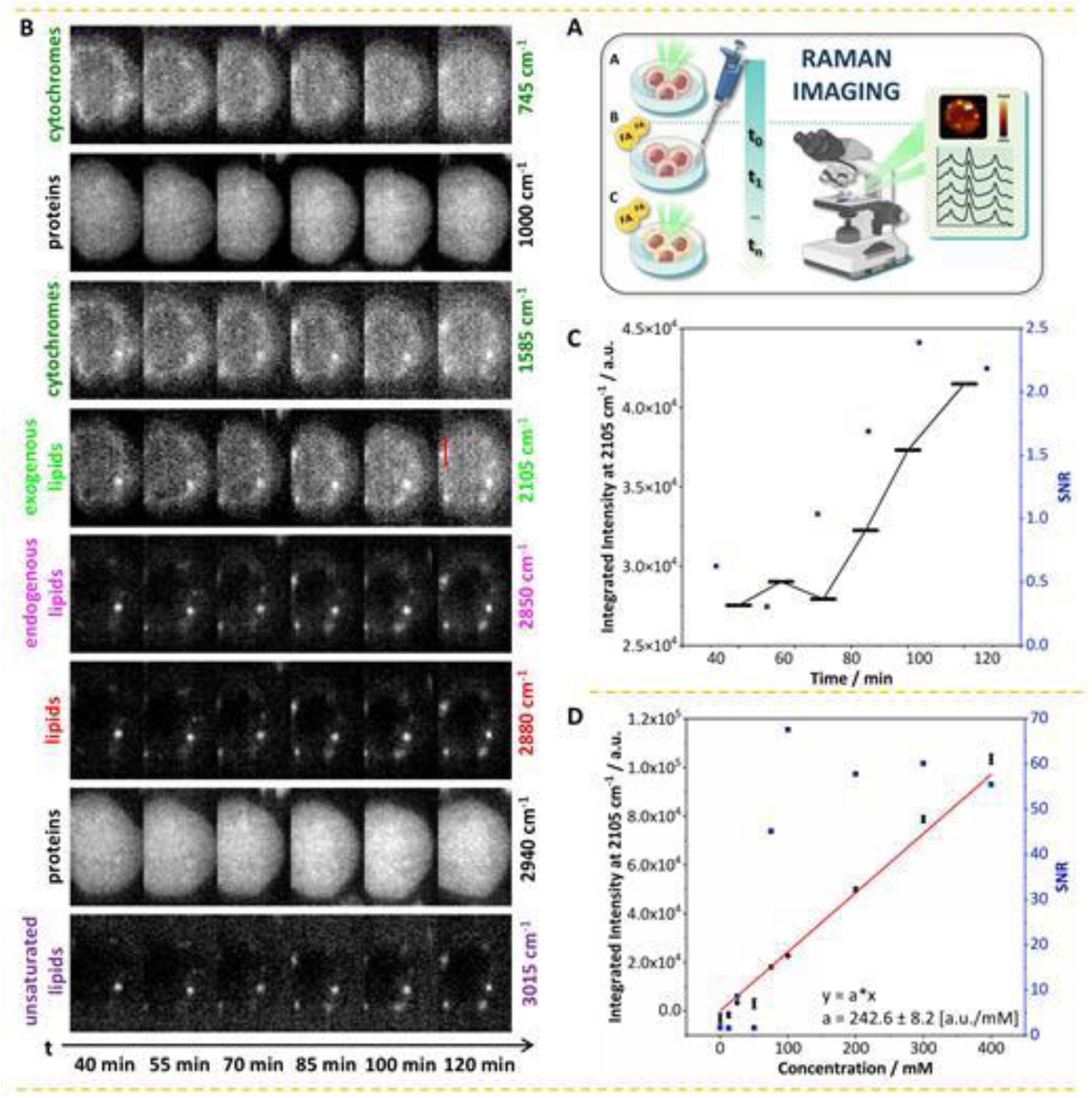
Real-time tracking of metabolic activity across multiple pathways at the subcellular level using RS. Schematic overview of experiments to track temporal metabolic transformations of dPA as a molecular isotopic tracer and Raman-based imaging (A). Spatiotemporal dynamics of selected biomolecules in a single live HL-60 cell, along with proteins (1000, 2936 cm^−1^), cytochromes (745 and 1585 cm^−1^), lipids (2880 cm^−1^), including exogenous lipids (2105 cm^−1^), endogenous lipids (2850 cm^−1^), and unsaturated lipids (3015 cm^−1^) (B). Time dependence of the total signal intensity of exogenous lipids analyzed on images presented in (B) (2105 cm^−1^) compared with SNR (C). SNR was evaluated along the intensity cross-section across the LD, marked in red. Calibration curve showing the relationship between signal intensity of dPA at 2105 cm^−1^ measured using RS and the concentration of deuterated fatty acid in DMSO.

The results presented in **Figure 2B** reveal the time course of exogenous PA accumulation in an exemplary live HL-60 cell incubated with dPA. Initially, dPA was incorporated into pre-existing LDs (exogenous and endogenous FA colocalize in the first tens of minutes of uptake), followed by the formation of new LDs that contain contributions from both exogenous and endogenous fatty acids. Furthermore, colocalization of cytochrome-associated proteins with LDs, including newly formed ones, was observed. Notably, both pre-existing and newly formed LDs exhibited partial unsaturation, indicating the presence of lipids containing C=C bonds. No significant alterations in cellular size or morphology were observed during the uptake process, as indicated by the consistent spatial distribution of total protein signals (1000, 2940 cm^−1^). However, a global decrease in the cytochrome-associated protein signal was observed due to photobleaching or cell stress (SI).

The presence of dPA–specific signals in the silent region of the Raman spectrum enabled semi-quantitative tracking of its uptake and accumulation over time by calculating the sum of the integrated signal intensity within the selected imaging plane. This temporal analysis, along with the corresponding signal-to-noise (SNR) assessment performed on images along a marked cross-section (red line, **Figure 2B**), is presented in **Figure 2C**. The initial constant signal level persisted for approximately 70 minutes, followed by a sharp rise that continued until the 120-minute time point. Although a temporal trend in signal intensity is evident, at this stage, without a quantitative analysis, it is not possible to accurately assess concentration changes over time or the kinetics of dPA uptake. As shown, during the RS measurements, the SNR was relatively low (around 3), which corresponds to the analytical detection limit for specific components. Additionally, the SNR was even lower for the first 70 minutes. This may explain why no increase in the signal was observed during the first 70 minutes. It seems that due to the high noise level, imaging based on RS is not the most efficient tool for detecting low analyte concentrations at the initial stages of the uptake. Nevertheless, it still enables monitoring of uptake processes in single cells and offers the vital advantage of simultaneously tracking multiple metabolic pathways in living cells.

The first step toward enabling kinetic studies, that is, analysis of rates of change in biochemical processes using RS, is the construction of a calibration curve to establish the relationship between signal intensity (in this case, dPA in the silent region of the Raman spectrum) and its concentration. This analysis was performed and is presented in **Figure 2D**. The resulting calibration curve shows a linear relationship between dPA signal intensity and concentration in DMSO. However, SNR analysis revealed limitations of the method, specifically, at concentrations below 100 mM, the Raman signal becomes increasingly obscured by noise, rendering quantitative analysis unreliable in this range. This limits our ability to track signal accumulation, particularly at the initial stages. It is also important to note that in RS-based measurements of single cells, the acquisition time for a hyperspectral map is equal to several minutes (8.25 min in this case). As a result, the recorded signal represents an accumulation over a defined time window rather than an instantaneous snapshot (marked as lines on the time-domain scale rather than single points, **Figure 2C**). Therefore, such measurements can be considered an integral analysis of kinetics, reflecting the average biochemical state over the duration of the acquisition, not capturing rapid, discrete temporal changes. Due to the long time required for recording signals, this type of measurement is challenging to perform for a large group of cells (in the presented example, it was a single exemplary cell), and therefore, it is difficult to assess cell-to-cell heterogeneity. However, on the other hand, it allows for the collection of a significant number of spectra (over 2,500 spectra collected from the cell at a single time point), providing comprehensive information on the spatiotemporal dynamics of metabolic transformations.

### 2.3. Rapid subcellular real-time tracking of dPA influx

The ability to acquire hyperspectral images using RS enables simultaneous tracking of multiple metabolic pathways (**Figure 2B**) and semi-quantitative tracking of dPA uptake. However, due to the inherently low SNR and the current limitation to integral kinetic measurements, this approach faces significant constraints in resolving the early stages of dFA uptake. A potential solution to these limitations is the use of SRS, which enables much faster point-wise measurements (~0.4 seconds/cell, **Figures 3 & 4**). This mode of signal acquisition can be referred to as differential kinetics analysis. **Figure 3A** presents the spatiotemporal dynamics of dPA uptake in three representative live cells (A) over a short time course (0 – 25 min) with very short time intervals (45 seconds). To capture such a short time regime and the dense sampling of dPA uptake, it was necessary to perform imaging in a single selected plane (2D), to use a higher dPA concentration (600 µM), and to use an SRS system with a 20 MHz repetition rate. The results are presented in **Figure 3**. On the other hand, the use of an 80 MHz laser system enabled the successful tracking of dPA accumulation throughout the entire cell volume, albeit under a different temporal regime, over longer time scales (5 – 60 min, **Figure 4**) and with extended intervals between consecutive measurements (~15 min). This experimental setup also allowed the use of a lower dPA concentration (200 µM), while maintaining sufficient signal quality for kinetic analysis. A 3D spatiotemporal analysis of subcellular uptake kinetics across a substantial population of cells (54 cells) is shown in **Figure 4**. With a 3D acquisition time of <1 second *per* cell, this approach maintains sufficient temporal resolution to be considered a differential kinetics analysis. However, compared to RS, it is not possible to obtain complete information about metabolic changes. Using SRS, it is possible to track the distribution of one type of molecule (exogenous FAs) over time, rather than the entire chemical fingerprint included in the RS spectra. This is due to the long time required to tune the laser wavelength in the SRS system.

**Figure 3.**
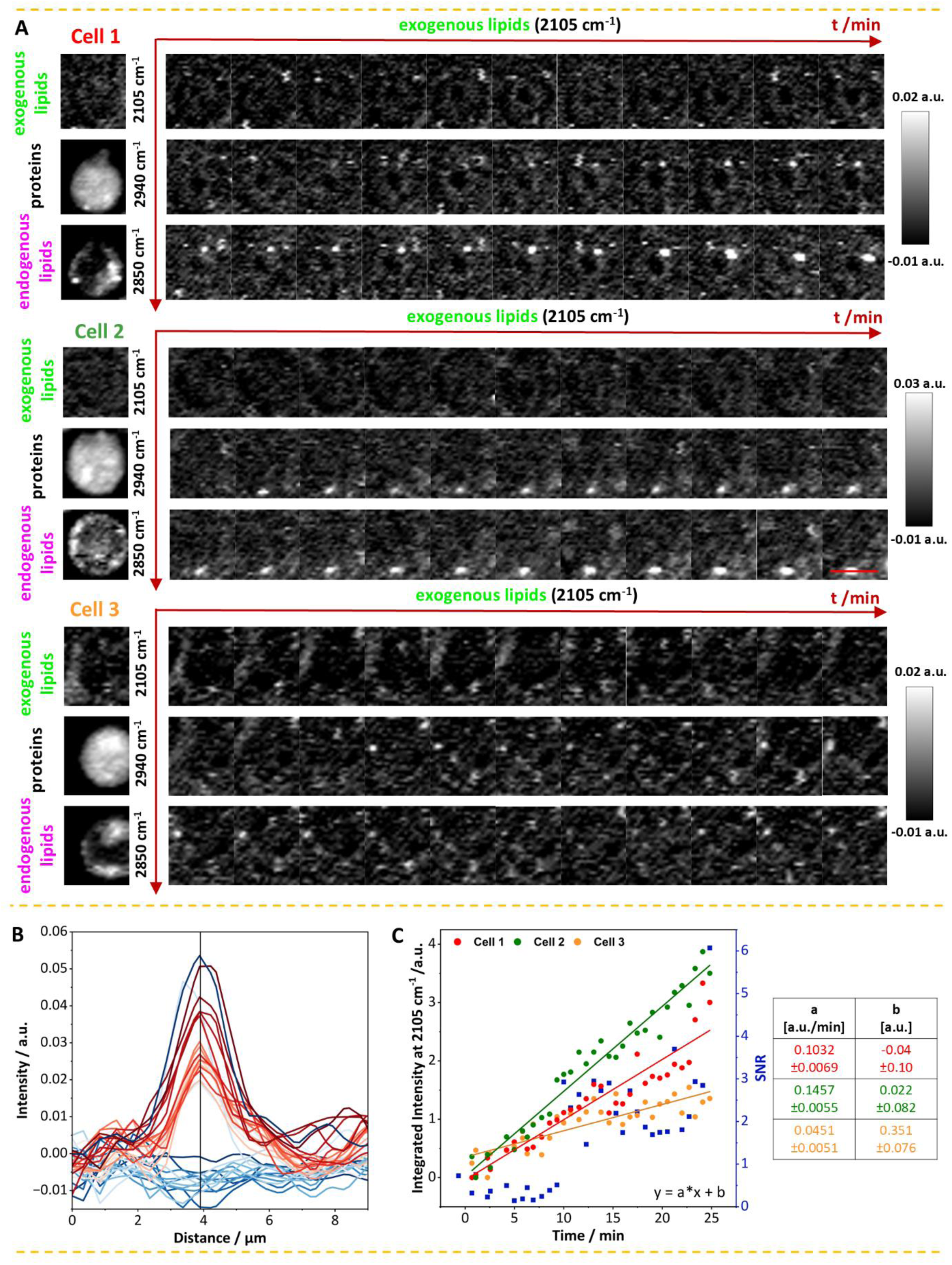
Time-resolved subcellular tracking of dPA uptake over a short time scale (0 – 25 min) using SRS. Spatiotemporal dynamics of dPA uptake in three representative live cells (A). For each cell, imaging was initially performed prior to dPA exposure to visualize cell morphology (proteins at 2940 cm^−1^), the distribution of endogenous lipids (2850 cm^−1^), and the accumulation of exogenous lipids (2105 cm^−1^). Following the addition of dPA, cells were imaged over a 0–25-minute time course to track dPA uptake (2105 cm^−1^). The acquisition time for a single map of a cell was ~0.4 seconds (A). Formation of a single LD visualized by time-resolved analysis of signal intensity along the cross-sectional profile of a selected LD in cell 2 (B). The cross-section was marked in red in the last image. Curves showing time dependence of the total signal intensity of exogenous lipids (2105 cm^−1^) for three cells presented in (A) compared with SNR calculated for cell no. 2 (C). Linear fit parameters for the individual kinetic curves are summarized in the table (C). R² values were equal to: 0.88, 0.96, and 0.72, respectively.

**Figure 4.**
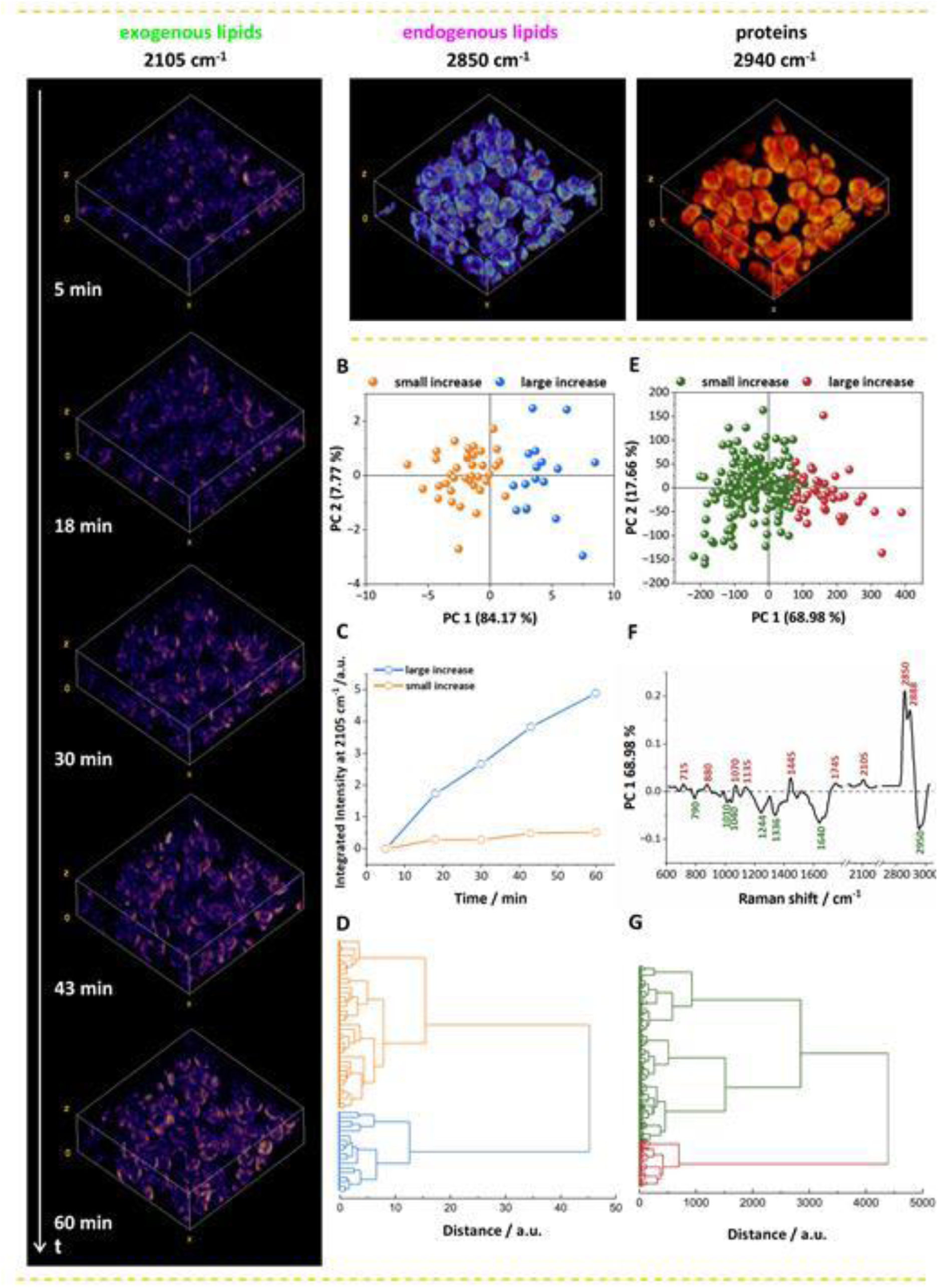
3D time-resolved subcellular tracking of dPA uptake over a longer time scale (5 – 60 min) using SRS. 3D spatiotemporal dynamics of dPA uptake in a representative cell population of 54 cells (A). SRS imaging was first performed prior to dPA exposure to visualize cell morphology (proteins, 2940 cm^−1^) and the distribution of endogenous lipids (2850 cm^−1^). Following the addition of dPA, cells were imaged over a 5–60-minute time course to track dPA uptake (2105 cm^−1^). The acquisition time for the entire 3D stack of one cell was approximately 0.4 seconds (A). Scores plot of PCA of curves showing time dependence of the total signal intensity for the whole cell volume of exogenous lipids (2105 cm^−1^, B) for the whole cell population presented in (A), together with mean curves calculated for groups identified by a cluster analysis (D). A dendrogram of a cluster analysis of time-dependent curves (D). The colors of the curves (B-D) correspond to the same groups identified by cluster analysis (D). Scores plot of PCA of cellular Raman spectra from fixed cells incubated with dPA for 45 minutes (E) together with a loading plot along the PC1 axis (F). A dendrogram of a cluster analysis of cellular Raman spectra analyzed in (E). The colors of the spectra and bands in (E-G) correspond to the same groups identified by cluster analysis (G).

For each cell shown in **Figure 3**, we analyzed cellular morphology (2940 cm^−1^) and the distribution of endogenous (2850 cm^−1^) and exogenous lipids (2105 cm^−1^) prior to dPA addition. Notably, all cells contained a baseline pool of endogenous lipids. Following the addition of dPA, its intracellular accumulation was tracked over a 0–25-minute time course. In cells 1 and 2, a clear accumulation of dPA was observed as single LDs, with a progressive increase in signal intensity over time. In contrast, cell 3 displayed a more diffuse cytoplasmic distribution of the dPA signal, with only small, sparsely formed LDs that were less pronounced compared to those in cells 1 and 2. This indicates cell-to-cell heterogeneity in dPA uptake and LD formation. As observed in the longer time-scale experiments shown in **Figure 2**, dPA accumulation was frequently detected in pre-existing lipid droplets; however, this was not always the case. In cell 2, for instance, we observed a gradual formation of new LDs over time, as evidenced by tracking dPA signal intensity along a defined cross-section of the droplet (**Figure 3B**).

The uptake and accumulation of dPA in single live cells were monitored semi-quantitatively over time by calculating the total dPA signal intensity within a selected imaging plane at each time point for individual cells. The results are presented as time-dependent uptake curves (**Figure 3C**). For cell no. 2, SNR was additionally calculated based on the signal intensity of exogenous lipids along a cross-section of a single LD (**Figure 3B**). As shown in **Figure 3B**, during the early phase of dPA uptake, a linear increase in signal intensity over time was observed, even in cell 3, where no distinct LD formation was detected. The uptake rates across all analyzed cells, calculated as the slopes of the fitted lines, varied slightly, showing differences in uptake between cells (**Figure 3C**). The findings further support the existence of significant cell-to-cell heterogeneity in FA uptake. A gradual increase in SNR was also observed, consistent with the accumulation of dPA signal over time. Notably, within the first 10 minutes, the signal intensity remained too low for reliable detection of the LD formation. A marked increase in SNR occurred only after this time point. Nevertheless, even at early points (<10 minutes), linear signal growth was still evident when integrating the total signal over the entire cell area rather than a single LD. Additionally, both signal intensity and SNR showed some variability across time points, likely due to slight movements of the non-adherent cells during measurement or fluctuations in laser power. Despite these limitations, our data demonstrates that dPA uptake proceeds at a steady rate during the initial minutes following addition, as evidenced by the linear time-dependent increase in dPA signal intensity (**Figure 3C**). Moreover, our results provide subcellular-level tracking of dPA uptake and allow monitoring of the dynamic increase in signal intensity at the level of individual LDs.

Figure 4A shows 3D time-resolved subcellular tracking of dPA uptake over a longer timescale (5 – 60 min) and with longer intervals (~15 minutes) than presented in Figure 3 but analyzed in the whole volume of cells and for a larger cell population (54 cells). Before the introduction of dPA, we assessed cell morphology (2940 cm^−1^) and endogenous lipid content (2850 cm^−1^) (Figure 4A). Once again, HL- 60 cells were found to contain pre-existing LDs, indicating an endogenous lipid reservoir prior to treatment.

A semi-quantitative analysis of dPA uptake was performed on individual cells across their entire volume. To achieve this, the total dPA signal was summed for each cell in 3D, and its intensity was tracked over time to generate time-dependent uptake curves. These temporal profiles were subsequently analyzed using PCA, with the results presented in Figure 4B. Figure 4C presents the average temporal profiles of two distinct cell groups identified by cluster analysis (Figure 4D). One group (~ 33% of cells) exhibited a pronounced and rapid increase in dPA signal intensity over time (described as ‘big increase’), and the other showed only a modest increase in signal intensity (described as ‘small increase’) (Figure 4C). It can also be observed that, even within the 5–60-minute uptake regime and across the entire cellular volume, the dPA signal increases steadily over time (Figure 4C). However, inspection of the PCA results reveals substantial dispersion along PC-1, indicating marked heterogeneity in dPA uptake among individual cells (Figure 4B). Capturing this variability was made possible by measuring a large cell population of 54 cells, which enabled the identification and characterization of single-cell uptake variability.

To better understand the observed differences in dPA uptake kinetics at the single-cell level and to explore whether these differences may be linked to cellular phenotype, we performed PCA analysis of RS spectra from fixed HL60 cells incubated with dPA for 45 minutes (217 cells, n = 3). The resulting PCA score plot (Figure 4E) revealed a similar degree of heterogeneity in uptake as previously observed in the kinetic analysis (Figure 4B). The groups of cellular spectra corresponding to high and low uptake profiles were identified through cluster analysis (Figure 4G). Notably, the proportion of cells displaying more substantial dPA uptake in the analyzed cell populations (21%, **Figure 4E&G**) closely matched the proportion of cells showing faster uptake kinetics in the live-cell time-course experiments (**Figure 4B&D**). Furthermore, the PCA loading plot (Figure 4F) highlights biochemical differences between cells exhibiting high versus low dPA signal intensity, suggesting divergent uptake behavior. Cells exhibiting higher FAs uptake showed not only an increased content of exogenous FAs, but also elevated levels of endogenous FAs (1445, 2850, 2880 cm^−1^), as well as triglycerides (1745 cm^−1^) and phospholipids (715, 1070 cm^−1^). In contrast, cells characterized by reduced FAs uptake exhibited an increased abundance of nucleic acids (790, 1244, 1336, 2950 cm^−1^) and protein content (1010, 1040, 1640, 2950 cm^−1^), indicating distinct metabolic profiles compared to high-uptake cells. A chemical composition similar to that of this subgroup of cells was observed in control cells. PCA conducted to compare control cells (236 cells, n = 3) with those incubated with dPA (217 cells, n = 3) confirmed that the observed biochemical differences in dPA-treated cells exhibiting lower dPA signals are related to reduced FA uptake. From a biochemical perspective, the composition of this group of cells is more like that of control cells. Overall, the presented analysis demonstrates that differences in dPA uptake kinetics are closely linked to the biochemical composition of cells. Furthermore, kinetic profiling not only captures cellular heterogeneity but may also serve as a distinguishing parameter for cell classification, analogous to chemical composition.

## 3. Discussion

The use of different spectroscopic imaging techniques (RS and SRS) enabled us to track dPA uptake by cells across various temporal regimes (from the first minutes up to 2 hours). By treating time as a unified, independent variable, results from different systems and laboratories were combined into an integrated temporal kinetic curve (Figure 5A). However, such integration requires a consistent definition of time progression across all experiments. In this study, the time point zero was defined as the moment of dPA addition to the cells. However, different measurement attempts conceptualized time differently. In SRS, where the acquisition time per cell is relatively short (~0.4 seconds), time was treated as a discrete point variable, enabling differential-kinetics analysis. In contrast, for RS, with longer acquisition durations, time was better represented as a continuous interval, aligning with an integral kinetic framework. Nevertheless, this didn’t prevent the formation of a unified kinetic curve. However, it is somewhat schematic, as each technique measures signal intensity *via* different physical effects. To quantitatively merge these datasets into a fully analyzable kinetic curve, it would be necessary to convert signal intensities into absolute concentrations for each experimental setup using well-standardized calibration curves. This would enable the construction of a complete, continuous kinetic profile. For example, we demonstrated that within the concentration range of up to 400 mM (Figure 2D), the Raman signal increases linearly with the dPA concentration. Overall, this study presents an initial step toward the goal of analyzing the kinetics of FA uptake analytically, demonstrating that the proposed methodological framework can generate a unified time-dependent curve over an extended temporal range that characterizes dPA uptake.

**Figure 5.**
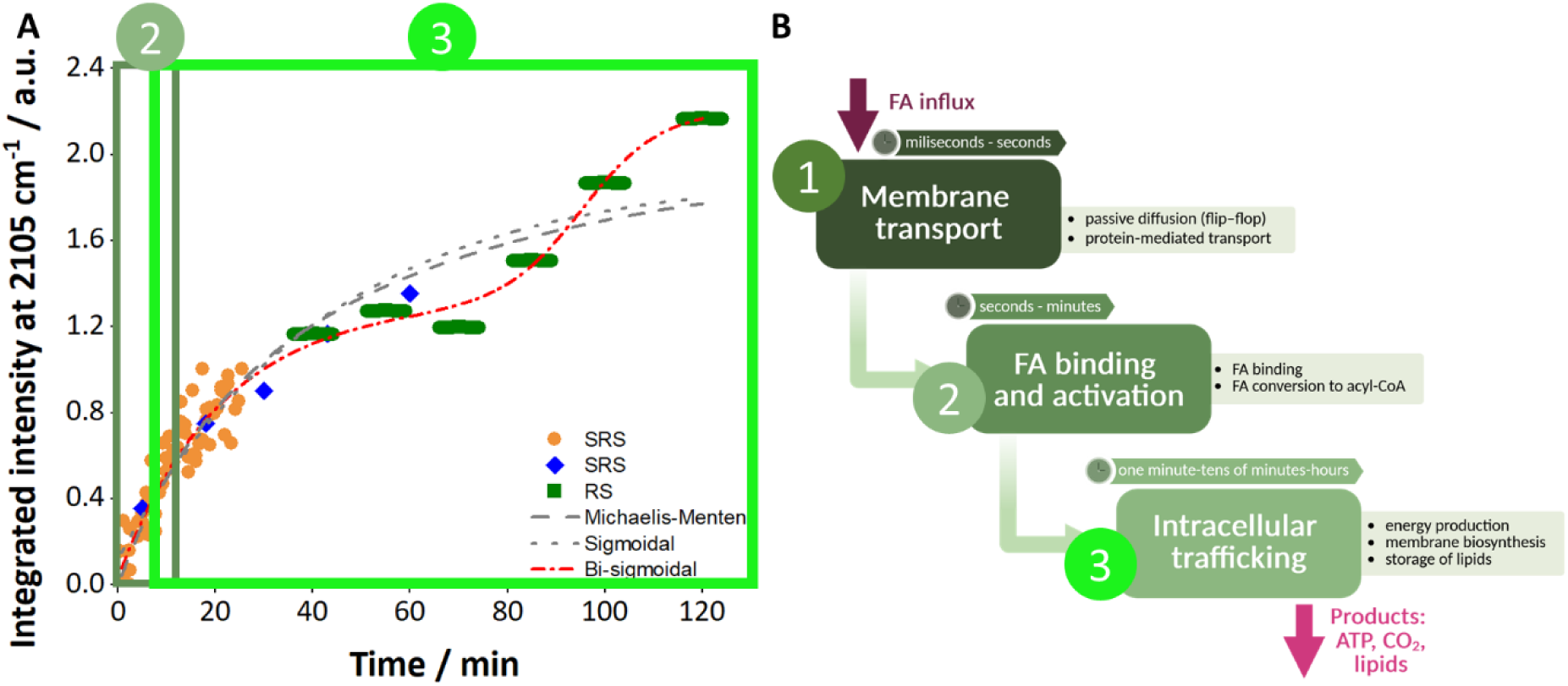
The integrated temporal curve of dPA uptake compared with the schematic representation of the sequential stages of fatty acid uptake and its initial metabolism. The integrated temporal curve of dPA uptake obtained using different methods (RS and SRS) and across different time regimes (from the first minutes to 2 hours) (A). Three different fits to the curve were also proposed: Michaelis-Menten, sigmoidal, and bi-sigmoidal. Three main stages of dPA uptake (B): membrane transport (1), FA binding and activation (2), and intracellular trafficking and metabolism (3). The respective stages of FA uptake and their associated time frames are color-coded on the integrated temporal curve (A).

A kinetic curve spanning a broad time range enabled mapping FA uptake kinetics to distinct intracellular processes (Figure 5B): (1) transport across the plasma membrane, (2) binding and activation of FA within the cytoplasm, and (3) intracellular trafficking and further metabolic processing ^[35,36]^. Each of these processes occurs on a unique timescale and has its circadian rhythm ^[37,38]^. Membrane transport is rapid, typically taking place within milliseconds to a few seconds ^[39–42]^. Next, the FAs undergo desorption in the cytoplasm, binding, and activation by being converted to acyl-CoA. These steps typically occur within a time frame ranging from a few seconds to several minutes ^[42–45]^. In the subsequent step, FAs are directed to specific organelles for further metabolism, such as energy production in mitochondria, membrane biosynthesis in the endoplasmic reticulum, or storage as LDs following esterification into triacylglycerols ^[6,35,36,46]^. This final stage occurs over a time scale ranging from several minutes to tens of minutes or even hours ^[47–49]^.

Correlating the uptake curve with cellular processes enables a model-based description of these phenomena *via* fitted curves that represent dPA transformations within cells. Despite lower temporal resolution at longer times (tens of seconds *vs* tens of minutes) and concentration differences in early measurements (600 µM for the first 25 minutes, 200 µM after the first 5 minutes), a clear upward trend in the intensity of the dPA signal is apparent. To describe subcellular processes mathematically, three different fitting models were tested: Michaelis-Menten (R² = 0.90), sigmoidal (R² = 0.89), and bi-sigmoidal (R² = 0.93). The bi-sigmoidal curve provided the best fit, reflecting the complex interplay of uptake and downstream metabolic reactions ^[35]^. Early linear phases likely reflect passive transport due to the high dPA concentration, whereas later deviations indicate active metabolic engagement. Approximations using segmental linear fitting or modified Michaelis–Menten kinetics effectively describe this behavior ^[33,50–52]^.

As demonstrated by our results, the spectroscopic techniques employed in this study do not allow for the resolution of all individual stages involved in FA uptake. For example, membrane transport cannot be directly analyzed because of its short time scale. The minimum acquisition time required to scan an entire cell (at least several tens of seconds) and the detection limits imposed, in part, by the relatively high noise levels inherent to Raman imaging, prevent Raman measurements on the timescale characteristic of acid passage through the membrane. Due to the complex structure of the cell, the most effective way to directly investigate FA transport across the plasma membrane is to use simplified membrane models, which allow real-time tracking of fatty acid penetration through the lipid bilayer ^[39,42,53]^. As an alternative, indirect approaches, such as selective silencing or inhibition of membrane-associated transport proteins, could help differentiate between passive and facilitated transport mechanisms ^[50,54–57]^. To access sub-microsecond dynamics, advanced ultrafast methods such as femtosecond SRS could provide real-time visualization of molecular events at the membrane interface ^[58,59]^. It is also worth noting that implementing the SRS system with a 20 MHz repetition rate was crucial for measuring uptake at its early stages, as it provides higher per-pulse energy than a laser with a higher repetition rate (i.e. 80 MHz for the other SRS system used in this study to measure uptake after 5 minutes) at the same average power. Using a relatively low repetition rate (20 MHz *vs* 80 Hz) enabled stronger instantaneous excitation per pulse while limiting average power, thereby reducing heating and photodamage.

In the SRS- and RS-based experiments, the observed processes likely reflect the binding and activation of FAs, which occur within seconds to minutes (Figure 5B). This is evidenced by the accumulation of FAs within the cytoplasm and the gradual increase in signal intensity during the early stages of FA uptake (**Figures 3 & 4**). Notably, we observe the intracellular trafficking of exogenous FA and its involvement in metabolism, together with dPA accumulation in both pre-existing and newly formed LDs. Additionally, colocalization with mitochondria suggests that the intracellular trafficking of dPA may also be linked to energy metabolism ^[46]^, consistent with FA transfer at droplet–mitochondria contact sites reported by Olzmann et al. ^[6,46]^. These findings underscore the importance of investigating intracellular lipid trafficking, a process that is increasingly recognized as a crucial aspect of lipid metabolism and signaling.

Until now, several attempts have been made to study the kinetics of FAs uptake using spectroscopic methods ^[30,33,60,61]^, yet the present study advanced this field by enabling the capture of the earliest uptake stages at intervals as short as 45 seconds. Through 3D uptake analysis of 54 cells, single-cell kinetics were statistically evaluated, providing new insights into cell-to-cell heterogeneity. Similar to the findings reported by Sibling *et al.,* the present study also revealed two significant cell populations that differed in their temporal profiles of FAs uptake curves ^[31]^. One group of cells exhibited strong dPA uptake, while the other, more numerous groups showed markedly lower uptake. In this work, however, the analysis was conducted on a larger cell population across the entire cell volume, providing a more robust statistical representation. Furthermore, Raman spectral analysis revealed that variations in uptake kinetics correlated with changes in the cells’ biochemical composition.

Our findings demonstrate that kinetic uptake profiling enriches classical spectroscopic analysis by adding a dynamic layer to cell characterization. Beyond revealing metabolic heterogeneity, this approach could enhance phenotyping precision and provide an early marker of pathological transformations. To validate this hypothesis, further systematic studies are required to enable fully quantitative kinetic analysis, including the use of appropriate calibration curves and reference standards. Such an approach would provide a robust framework for assessing uptake kinetics and may ultimately offer a novel diagnostic dimension for applications in personalized medicine.

## 4. Conclusions

The development of many lifestyle-related diseases is closely associated with alterations in cellular lipid composition. Current diagnostics usually assess biomolecule levels at a single time point, yet growing evidence suggests that monitoring lipid metabolism dynamics, accounting for subcellular processes and single-cell kinetics, may yield a more precise diagnostic marker, particularly for detecting early stages of disease development.

This study expands current lipid kinetics research, primarily by filling the existing gap in understanding the early stages of FA uptake and extending analysis to a statistically significant cell population. Uptake of a labeled PA was monitored over different temporal regimes, from the initial minutes of uptake to 2 hours, using SRS and RS imaging. Rapid SRS imaging (0.4 s per cell) enabled differential kinetic analysis at high temporal resolution, while RS imaging (8.25 s per cell) captured integrational kinetics over longer timescales. Combining both approaches provided a continuous description of FA uptake, fitted with a sigmoidal curve that reflects complex lipid metabolism dynamics. Additionally, RS was used to perform simultaneous spatiotemporal analysis of a few biomolecules, revealing that during dPA uptake, they were first incorporated into pre-existing LDs, followed by the formation of new, partially unsaturated LDs associated with cytochrome-related proteins.

A 3D imaging across a large cell population provided statistical evaluation of single-cell kinetics, showing pronounced cell-to-cell heterogeneity in dPA uptake dynamics. Two subpopulations were identified: cells with strong uptake activity and a larger group with considerably lower activity. Raman spectral data linked these kinetic differences to distinct biochemical composition. Moreover, heterogeneous accumulation of dPA within individual LDs demonstrated that lipid metabolism is heterogeneous not only between cells but also within single cells at the subcellular level.

Despite certain limitations in detection, spectroscopic approaches proved effective for monitoring deuterated palmitic acid uptake and capturing heterogeneous metabolic behavior. This heterogeneity could become a valuable parameter for early detection of metabolic disturbances at the single-cell level. Future work should focus on standardizing measurement protocols and developing quantitative methods for kinetic modeling, enabling comprehensive lipid flux characterization in living cells.

## 5. Material and methods

### 5.1 Incubation of HL-60 cells with FA

HL-60 cells (human promyelocytic leukaemia) were cultured in RPMI 1640 medium (Roswell Park Memorial Institute, Thermo Fisher) with 10% FBS (Fetal Bovine Serum, Gibco) under 5% CO_2_ and 37^0^C (ESCO Cell culture incubator). For experiments, 1×10^6^ cells/ml were seeded in a 6-well plate and incubated for 45min with 200 μM: PA/dPA. Before supplementation, FAs/dFAs were saponified with NaOH ^[62]^ and conjugated to 10% FA-free BSA (bovine serum albumin, Sigma Aldrich). Control cells received BSA alone. After incubation cells were washed with warm HBSS (Hanks’ Balanced Salt solution, Gibco) and twice with warm PBS (Phosphate Buffer Saline, Gibco). For statistical measurements, cells were fixed with 0.5% glutaraldehyde (Sigma-Aldrich) for 10 minutes at room temperature and washed three times with PBS to remove excess fixative. The prepared cells were resuspended in PBS and kept at 4^0^C until Raman measurements. Live-cell measurements were performed up to 45 min in warm solution. FL (fluorescence) microscopy was used to analyse FAs/dFAs cytotoxicity and LDs distribution.

### 5.2. Spontaneous Raman imaging of HL-60 cells

Raman measurements of fixed HL-60 cells and chemometric analysis was done. For live cell measurements, HL-60 cells (5×10^5^ cells/ml) were seeded in 35 mm dishes with 12 mm diameter quartz-bottom substrates (SF-S-D12, Fine Plus International). RS imaging of live cells were conducted using a homemade slit-scanning Raman microscope, equipped with 532 nm laser generator (Millennia eV, Spectra-Physics) and 60×/1.27NA water immersion lens (CFI Plan Apo IR 60XC WI, Nikon) with a laser power density of 1.5 mW/μm^2^. The system acquired up to 2048 spectra simultaneously with ~4 cm^−1^ spectral resolution and an exposure time of 3 s/line. For each measurement, an area was scanned by 45 lines with 200 spectra per line at an approximate pitch of 0.25 μm, enabling the collection of 9,000 spectra in a single measurement (8.25 min/area).

### 5.4. Stimulated Raman imaging of HL-60 cells

Stimulated Raman imaging was conducted using two different home-built SRS microscopes. The first system (Jagiellonian University in Krakow) ^[63–65]^ employed picosecond pulsed laser (2 ps, 20 MHz; Lazurite, Fluence) generating pump (750–950 nm, ~100 mW) and Stokes (1029 nm, ~450 mW) beams. The Stokes beam was modulated at 4 MHz (acousto-optic modulator) and combined with the pump before an inverted microscope (Ti2, Nikon) equipped with galvanometric scanning. The sample was illuminated through a 40×/0.95NA air objective (UPLXAPO40X, Olympus), while the scattered light was collected with a 50×/1NA water-dipping objective (MRD07620, Nikon). Detection was achieved through lock-in amplification and digitization via WITec system (WITec software and hardware). The imaging parameters was set at: pixel dwell time of 900 µs/pixel, a lateral pixel size of 500 nm (x, y), average imaging time per cell was 0.4 s and 45s of temporal intervals for analysis of the uptake.

In the second SRS system, a mode-locked Yb-fiber laser (Emerald Engine, APE) provided an output 2-ps laser pulse at 1031 nm with a repetition rate of 80 MHz, which served as the Stokes beam for SRS imaging. An optical parametric oscillator was used to tune the wavelength of the pump beam to obtain SRS signals at chosen Raman bands. The two laser pulses were focused on the samples by a water immersion objective lens (Plan Apo IR 60x, 1.27 NA, Nikon) and collected by another identical objective lens. Laser scanning was performed with a pair of galvanometer mirrors (VM500+, General Scanning Inc.). Two identical bandpass filters (FF01-850/310-25, Semrock) were employed to block the Stokes beam, and the pump beam was detected by a large-area silicon photodiode (PD, S3590-09, Hamamatsu Photonics). The signal from PD was filtered with a bandpass filter (BBP-21.4+, Mini-Circuits) to reject DC and high-frequency signals originating from laser repetition. The signal was amplified by a pre-amplified with a gain of 46 dB (SA-230F5, NF Corporation) and sent to a lock-in amplifier (UHFLI, Zurich Instruments) for demodulation. We measured Raman shifts at 3015, 2940, and 2850 cm^−1^ with a pixel dwell time of 250 µs and average powers of 75 and 150 mW for the pump and Stokes beam, respectively. To detect 2105 cm^−1^, we increased the average powers to 130 and 260 mW of the pump and Stokes beam, respectively, to detect relatively weaker signals. 3D SRS imaging was performed layer-by-layer in the z-direction, from the top to the bottom of the cell, using a piezo scanner. The imaging parameters included a pixel dwell time of 50 µs/pixel, a lateral pixel size of 434.78 nm (x, y), and a z-step spacing of 1 µm. The average imaging time per cell was 0.4 s. The temporal intervals for analysis of the uptake were equal to 15 minutes.

The kinetic curves for the 2D SRS images were obtained by summing the signal for each imaging plane over the entire cell and plotting its dependence on time. To construct kinetic curves from the 3D SRS images, the signal corresponding to exogenous fatty acids (2105 cm^−1^) was summed for all imaging planes for each cell separately. The resulting integrated signal was then plotted as a function of time. The kinetic curves obtained for individual cells were subsequently subjected to chemometric analysis, including cluster analysis and principal component analysis (PCA).

## Abbreviations

FAs: fatty acids
dFAs: deuterated fatty acids
HL-60: human promyelocytic leukaemia
LDs: lipid droplets
PA: palmitic acid
dPA: palmitic-d_31_ acid
PCA: principal component analysis
RS: spontaneous Raman spectroscopy
SNR: signal-to-noise ratio
SRS: stimulated Raman spectroscopy

## Acknowledgements

This work was partially supported by „Label-free and rapid optical imaging, detection and sorting of leukemia cells” project, which is carried out within the Team-Net program (POIR.04.04.00-00-16ED/18-00) of the Foundation for Polish Science co-financed by the European Union under the European Regional Development Fund. This research was funded in part by the National Science Center Poland (NCN), Miniatura (2023/07/X/ST4/01573) to AMN and OPUS (2024/53/B/ST4/02698) to MB, and by a grant from the Faculty of Chemistry under the Strategic Programme Excellence Initiative at Jagiellonian University to AMN. For Open Access, the author has applied a CC-BY public copyright license to any Author Accepted Manuscript (AAM) version arising from this submission. ABD acknowledges support by the Foundation for Polish Science (Start 006.2025). Special acknowledgments are addressed to Atsushi Nakayama for helping to organize SRS measurements and Adriana Adamczyk for technical support.

## Availability of data and material

The datasets generated during and/or analysed during the current study are available from the corresponding author upon reasonable request.

## Authors’ contributions

Investigation – A.B-D., P.D., T.K., X.H. I.Y., M.L. J.Q.; Formal analysis – A.B-D., P.D., X.H. (Raman imaging of live cells), I.Y. (SRS imaging of living cells), M.C. (Fluorescence imaging), K.S. (Raman imaging of fixed cells); Methodology – A.B-D., A.M.N.; Visualization – A.B-D., P.D.; Writing - original draft – A.B-D., P.D., T.K., X.H., I.Y., M.L.; Supervision – K.F., M.B., J.P-P, K.M.; Resources – K.F., M.B., K.M.; Validation – K.F., M.B., K.M., A.M.N., J.P-P; Writing - review & editing – A.B-D., K.F., M.B., J.P-P, K.M., A.M.N.; Funding acquisition – M.B., K.M, A.M.N.; Conceptualization – A.M.N.

## Ethics approval

All procedures performed in studies involving human leukemic cells, obtained as a commercially available *in vitro* model, were in accordance with the ethical standards in the World Medical Association

